# Phytochrome mediated responses in *Agrobacterium fabrum*: growth, swimming, plant infection and interbacterial competition

**DOI:** 10.1101/2020.04.24.060905

**Authors:** Peng Xue, Yingnan Bai, Gregor Rottwinkel, Elizaveta Averbukh, Yuanyuan Ma, Thomas Roeder, Patrick Scheerer, Norbert Krauß, Tilman Lamparter

## Abstract

The soil bacterium *Agrobacterium fabrum* C58 infects plants by a unique DNA transfer mechanism. *A. fabrum* has two phytochrome photoreceptors, Agp1 and Agp2. We found that DNA transfer into plants by *A. fabrum* is down regulated by light and that phytochrome knockout mutants have diminished DNA transfer rates. The regulation pattern matches with that of bacterial conjugation reported earlier. Growth, swimming and interbacterial competition were also affected in phytochrome knockout mutants, although these effects were often not affected by light. We can thus distinguish between light-regulated and light-independent phytochrome responses. In microarray studies, transcription of only 4 genes was affected by light, indicating that most light responses are regulated post-transcriptionally. In a mass spectrometery-based proteomic study, 24 proteins were different between light and dark grown bacteria, whereas 382 proteins differed between wild type and phytochrome knockout mutants, pointing again to light-dependent and light-independent roles of Agp1 and Agp2.

## Introduction

Soil bacteria of the genus *Agrobacterium* can transfer genes into plants and thereby induce the formation of plant tumors. This infection causes massive losses in agriculture, but the mechanism is on the other hand used for plant transformation in many research laboratories (1). Do these soil bacteria respond to light, despite the anticipated dark environment they envisage? The uppermost part of the soil is not completely dark, as sunlight penetrates it to several mm depth, and plant roots guide red light several cm into the soil. In addition, *Agrobacterium fabrum* cells were also found on plant stems and leaves, *i.e.* in open sunlight environment (2). The sequencing of *A. fabrum* C58 revealed two phytochrome photoreceptors, termed Agp1 and Agp2 (3). Phytochromes sense light in the blue, red and far-red range of the visible spectrum. A large number of developmental effects are controlled by phytochromes in land plants (4) and fungi (5), and several examples of phytochrome effects in bacteria have been reported, including control of photosynthesis and infection of plants by plant pathogens (6-12). In a search for phytochrome responses of *A. fabrum*, however, knockout mutants were generated which had no apparent phenotype (13). A first clear *Agrobacterium* phytochrome response was found by a codistribution study (14, 15). In this investigation, Agp1 and Agp2 BLAST homologs were found in an almost identical subset of Rhizobiales species as BLAST homologs of TraA, a central player in bacterial conjugation. Experimental tests showed that conjugation of *A. fabrum* is indeed regulated by light and phytochrome. When strains with Ti plasmid were used, conjugation was drastically reduced by red or far-red and in *agp1*^-^ or *agp2*^*-*^ knockout mutants. In the *agp1*^*-*^ */ agp2*^*-*^ double knockout, no conjugation was observed.

In the present work, we tested the role of Agp1/2 on plant infection. Light effects on plant transformation have been reported several times, but the observations were not consistent. In some cases light increased transformation rates, in other cases it resulted in a reduction (16, 17), It is also not known whether photoreceptors of the plant or of *Agrobacterium* are responsible for the light regulation. The infection process of *Agrobacterium* species is coupled to a gene transfer which results in a stable integration of the “T-DNA” into the plant genome (18-20). This process and the process of conjugation have some common features. In both cases, a single stranded DNA is generated, induced by a nick at one or two positions on the plasmid. In case of infection, the mobile DNA (T-DNA) is a part of the tumor inducing plasmid, Ti-plasmid, and contains genes for phytohormone synthesis and production of amino acid derivatives. In both processes, single stranded DNA is covalently attached to the protein that makes the nick, which is VirD2 (21) in *A. fabrum* plant infection and TraA (22) in conjugation. During conjugation, the entire plasmid is transferred. The complex of protein and single-strand DNA is transferred through pili formed by the type IV secretion system into the target cells.

We also tested the impact of phytochromes in *A. fabrum* on growth, motility, and interbacterial competition and also performed microarray and proteome studies. These data show that protein levels are affected for few gene products and suggests regulatory mechanisms independent on transcription or translation. On the organismal level and on the level of single proteins, we found phytochrome effects that are light dependent and such that are light independent.

## Results

### Distribution of phytochromes in *Agrobacterium* strains

The present studies were performed with *Agrobacterium fabrum* C58, a strain which is often used as model organism and the *Agrobacterium* species which was sequenced first. Meanwhile, the genomes of several members of the genus *Agrobacterium* are available.

The number of phytochrome genes within an organism varies and the distribution of phytochromes among related species can provide information about their functions (23). We have therefore searched for homologs of *A. fabrum* phytochromes Agp1 and Agp2 in 21 other *Agrobacterium* species. All strains except one contain an Agp2 homolog, whereas only four (including *Agrobacterium fabrum* C58) contain an Agp1 homolog, and there was no *Agrobacterium* species without a phytochrome (***Figure 1***), suggesting that all strains can respond to light. Agp2 of *A. fabrum* and related phytochromes from *Rhizobiaceae* (15, 24, 25) belong to the bathy phytochromes which have a Pfr dark state and absorb maximally in the 750 nm wavelength range. Since among the Agp2 homologs in the *Agrobacterium* strains, all amino acids in the chromophore pocket (26) are conserved, we predict that these are all bathy phytochromes. This could indicate that detection of the long wavelength red light around 750 nm is most important for the fitness of *Agrobacterium* species.

**Figure 1.**
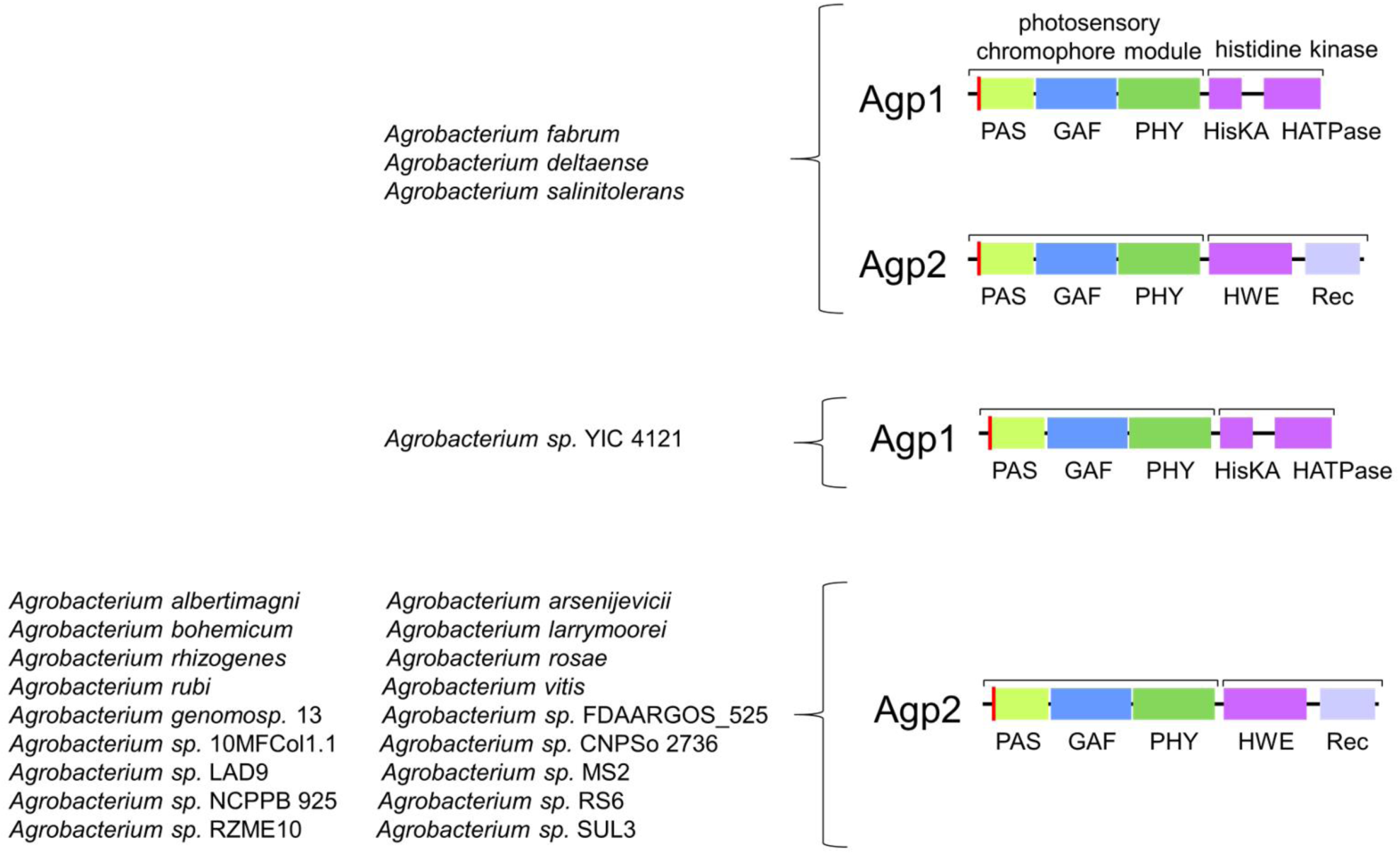
Distribution of Agp1 and Agp2 homologs in species of the genus *Agrobacterium*. Genome sequences are from NCBI, some genomes are published in (43-52).

### Growth in liquid medium

In these studies we measured the increase of cell densities of wild type *A. fabrum*, the *agp1*^*-*^, *agp2*^*-*^ knockout mutants and the *agp1*^*-*^ */ agp2*^*-*^ double knockout mutant in liquid cultures. In earlier work we found no effect of phytochromes on *A. fabrum* growth during 6 h (13). In the present study, we expanded the growth period to 51 h and performed assays at different temperatures, because Agp1 might also function as a thermosensor (15, 24, 27). Up to cultivation times of 6 h, the growth at ambient temperature was again not significantly different between wild type and mutants, but during prolonged growth times, all three knockout mutants displayed faster growth than the wild type (***Figure 2A-B***). The *agp1*^*-*^ knockout strain had the fastest growth and reached 2x higher cell densities than the wild type. Despite the clear mutant effects, we observed no impact of light on the growth (***Figure 2B***). This phytochrome effect at ambient temperature may be classified as “light-independent” (see below). At 37 °C, the cell densities of the *agp2*^*-*^ and *agp1*^*-*^ */ agp2*^*-*^ mutants if grown in the dark were lower as compared to the wild type and there was a clear inhibitory effect of red light on wild type and the *agp1*^*-*^ mutant. Therefore, red light regulation at 37 °C is mediated through Agp2. This effect is thus classified as “light dependent”. White light resulted in a slight increase of growth in wild type and *agp1*^*-*^ knockout and a strong increase in the *agp2*^*-*^ knockout and – remarkably – an induction of growth was induced by white light in the double knockout (***Figure 2***). *A. fabrum* has no LOV or BLUF protein homolog, which could serve as alternative photoreceptors. The photolyase PhrB, which has been identified by a blue light effect on motility (28, 29) is probably the photoreceptor of these light responses in the absence of phytochromes.

**Figure 2.**
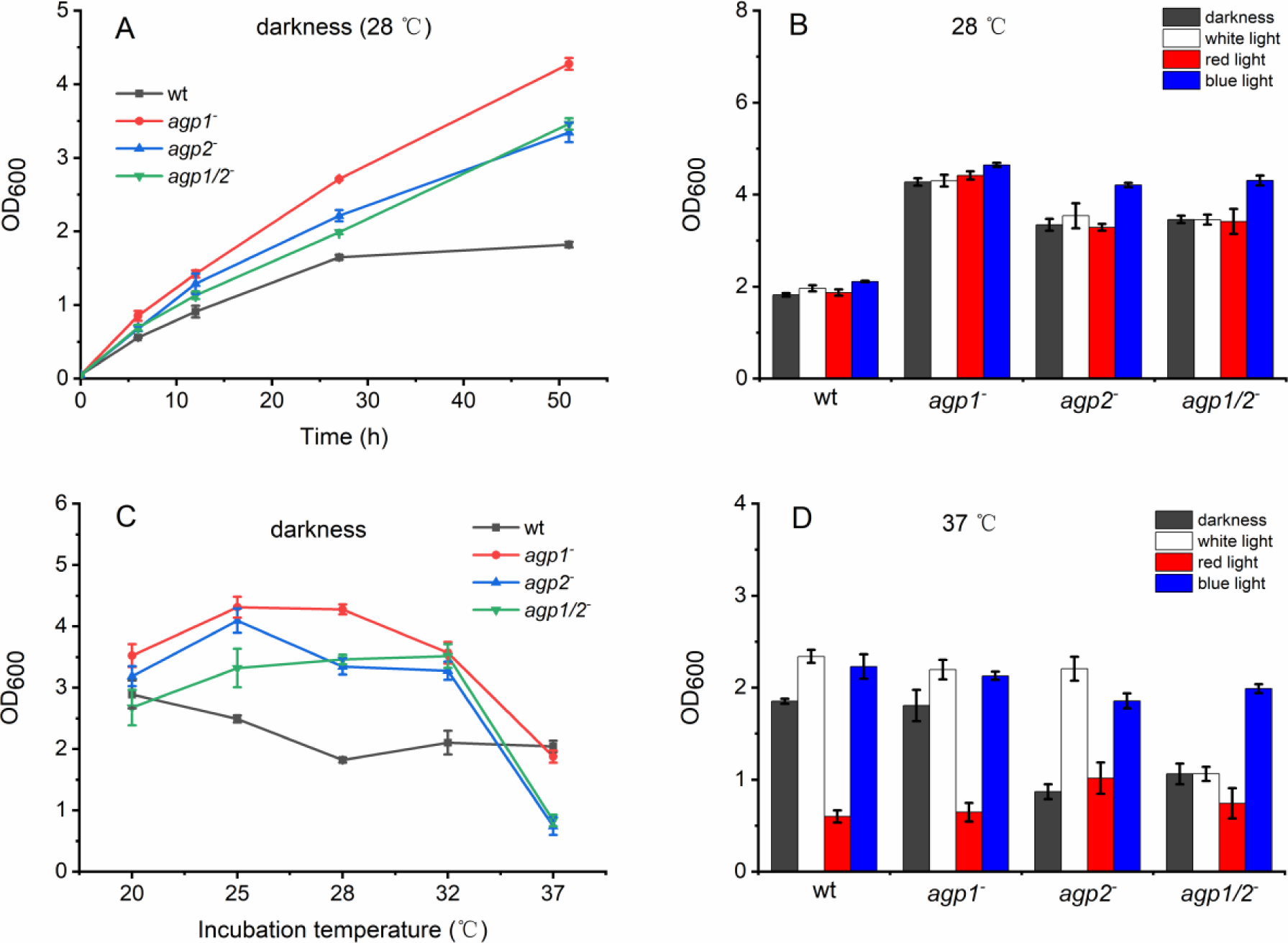
Effects of phytochromes Agp1 and Agp2 on growth of *A. fabrum* as measured by the optical density at 600 nm. Time curves of WT and mutants of *A. fabrum* in darkness at 28 °C (A) and a comparison of their growth under different temperatures after 51 h incubation (C). Effects of white, red and blue light on cell density 51 h after inoculation, cultivation at 28 °C (B) and 37 °C (D). Light intensities were always 40 µmol m^-2^ s^-1^. Mean values of three independent experiments ± SE.

### Cell motility

In earlier swimming plate studies light reduced the motility of *A. fabrum* significantly; this effect led to the discovery of the (6-4) photolyase PhrB (30). We performed here similar studies with the focus on phytochrome effects. The assays were performed at different pH values because spectral properties of Agp2 are pH dependent (31) (***Figure 3*** and ***Figure 4***). As above, the assays were performed at two different temperatures (here 26 °C and 37 °C). Under most conditions, colony diameters of *A. fabrum* were smaller at pH 5 or pH 9 as compared to the neutral range of pH 6-8. Exceptions for pH 5 are wild type in Figs 4a and 4d. In the neutral range, the single knockout mutants had slightly larger diameters than the wild type, whereas diameters of the *agp1*^-^ */ agp2*^-^ double knockout were slightly smaller (e.g. ***Figure 4A***). The pH -, mutant - and light - dependent pattern of motility is complex and does not allow to identify clear light or phytochrome effects. However, when dark minus light differences were calculated, a clearer pattern emerged.

**Figure 3.**
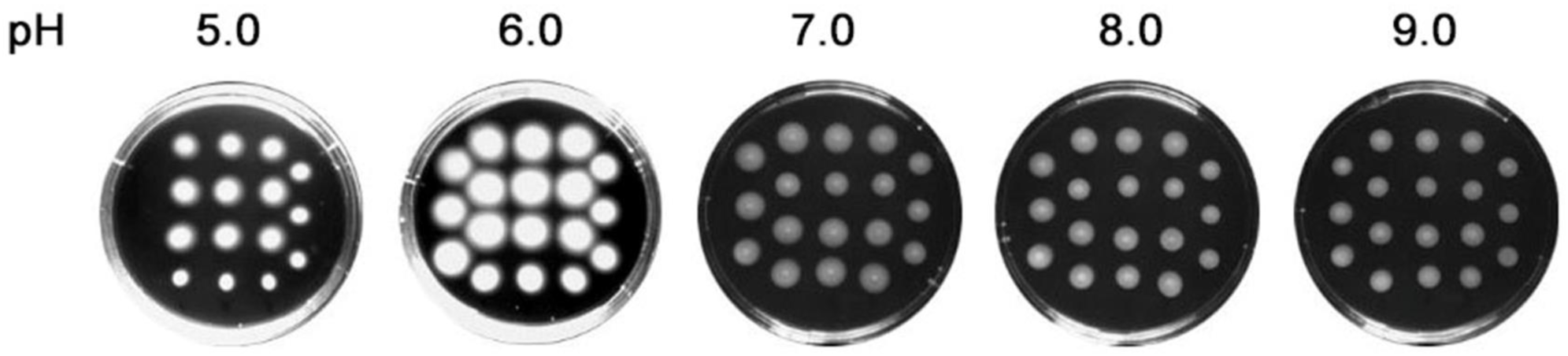
Results of *A. fabrum* swimming assay; wild type on 0.5% LB solid medium under different pH conditions.

**Figure 4.**
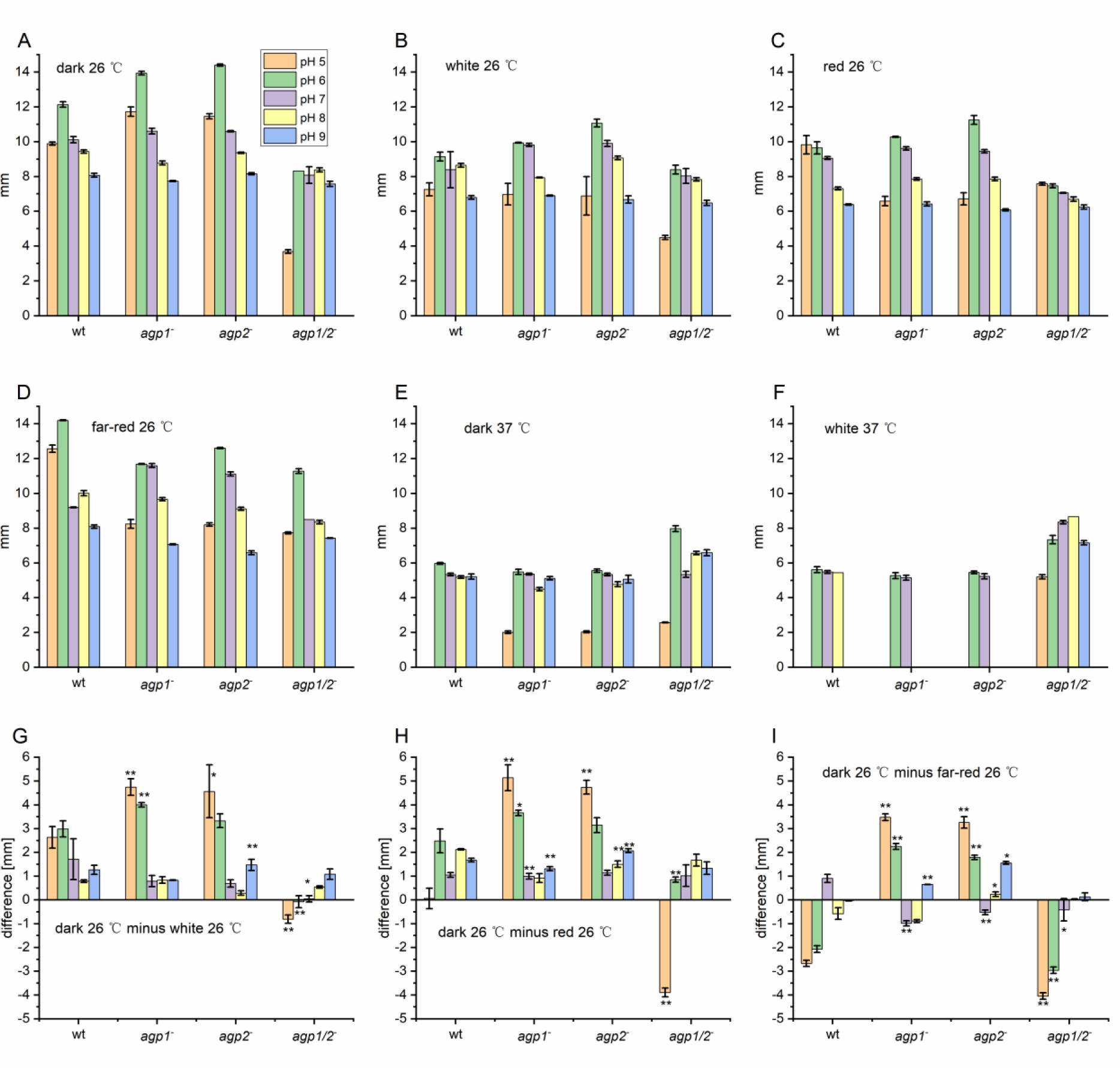
Cell motility of *A. fabrum*. The colony diameters were determined 30 h after inoculation at the indicated temperature and light conditions. Light intensities were always 40 µmol m^-2^ s^-1^. On each plate, 18 colonies were inoculated under identical conditions and mean values calculated. For the difference (panels G, H, I), values from light irradiatiated samples (as in panels B,C,D) were subtracted from dark samples (as in panel A). Three independent experiments with 18 colonies for each treatment, mean values ± SE.

These differences between dark- and light grown cultures are presented in ***Figure 4G-I***. In almost all cases was the difference at pH 5 the largest as compared to results at other pH values of the same treatment. We therefore concentrate on the pH 5 results in the following. For the wild type the difference was moderately positive in white-illuminated samples (***Figure 4G***), zero in red-illuminated samples (***Figure 4H***) and negative in far red-illuminated samples. These results show that there is an impact of light on motility. The single knockout mutants showed a positive difference of 3.5 mm - 5 mm in white, red and far-red light. The values were larger than for wild type or double knockout. These results indicate that both phytochromes mediate a light effect on motility but at the same time inhibit the action of the respective other phytochrome in the wild type. Therefore, a single knockout uncovers the light effect. In several conjugation experiments (experiments without Ti-plasmid and mutants in donor strain) phytochrome knockout also lead to enhanced light effect (15), and the stronger response in a single knockout as compared to the wild type seen there is reminiscent of the results from the growth experiments above.

Under far-red, wild type and double knockout showed negative differences in colony diameters (***Figure 4I***), just opposite to the general pattern. How could a far-red light effect be mediated in the double knockout without phytochrome? We consider an effect of unavoidable subtle temperature increase.

The 37 °C experiments were performed in dark and white light only. White light induced cell death at pH 9 in wild type cells and at pH 9, pH 8 and pH 5 in single knockout mutants (***Figure 4F-G***), as seen by the zero diameters under these conditions. In the double knockout, no cell death was induced by white light. This result shows that the effect of light on cell death is mediated through phytochrome(s) in wild type and single knockouts.

### Plant infection

Since conjugation of *A. fabrum* is under control of Agp1 and Agp2 (15), and plant virulence of other bacteria such as *Pseudomonas syringae* are phytochrome controlled (11), we investigated the effect of phytochromes on plant tumor induction by *A. fabrum* in several assays.

In a root infection assay, root segments of young *Arabidopsis thalina* plants were co-cultivated with *A. fabrum* cell cultures and the number of root segments that formed a tumor were counted (***Figure 5B***). Reproducibility of these assays was dependent on controlled temperature and light conditions during plant growth and precise cutting of the root segments. We therefore concentrated on a few experiments under controlled conditions. The proportion of root segments of *A. thaliana* in which *A. fabrum* induced tumors was almost 30% under these test conditions in the dark. This was down regulated in red light to 8%, which is significantly different from the dark value. With the double knockout mutant of *A. fabrum* in only 3% and 1% of the root segments tumors were formed in the dark and in red light, respectively (***Figure 5A***). The latter dark minus red difference was not significant. That in the double knockout mutant almost no tumor formation was observed suggests that the light inhibition of tumor formation is regulated through the *A. fabrum* phytochromes and that the plant phytochrome, which could be present in the root, is not relevant for the light regulation.

**Figure 5.**
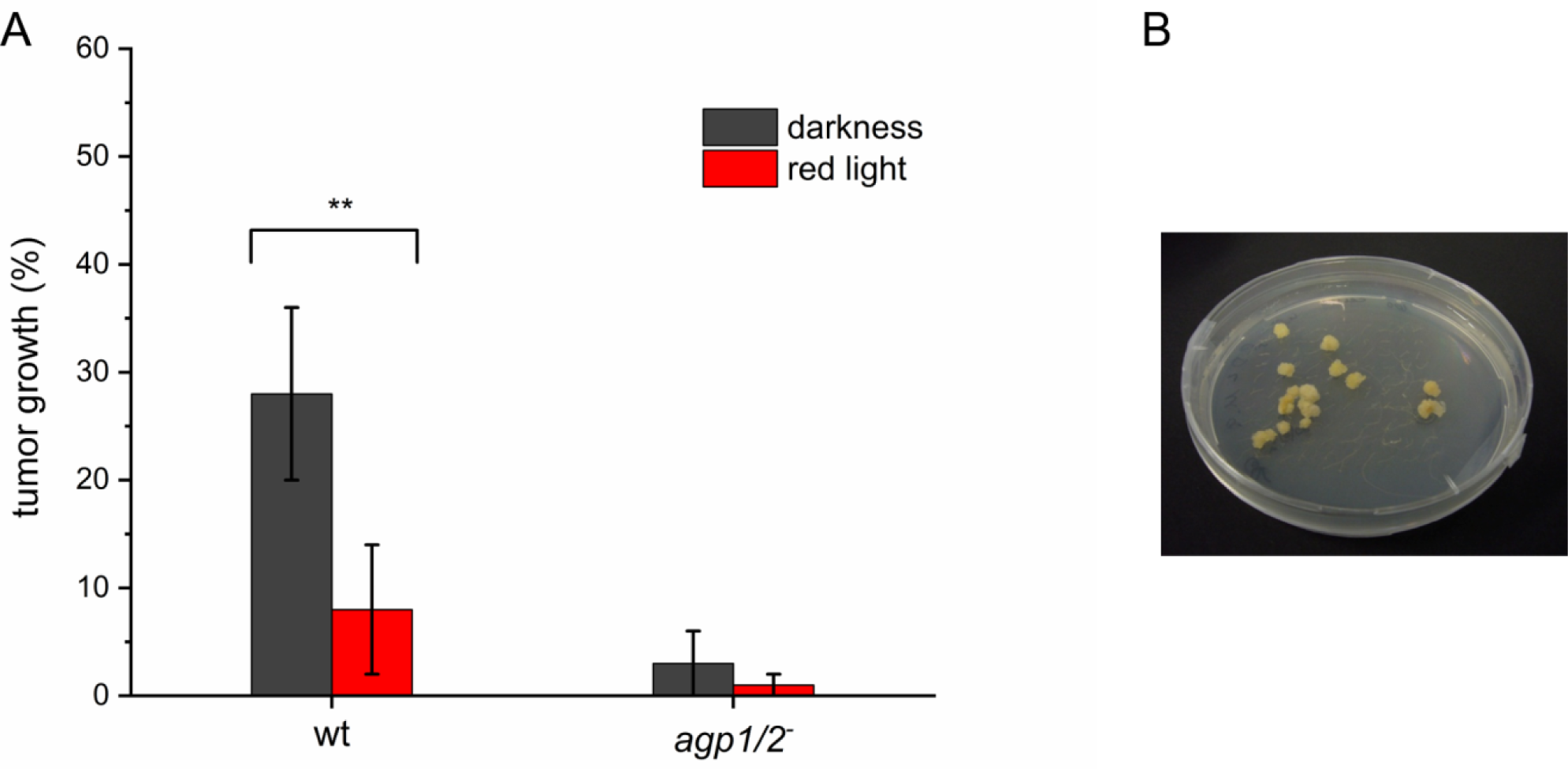
(A) *Arabidopsis thaliana* root infection by *A. fabrum* wild type and *agp1*^-^ / *agp2*^-^ double knockout mutant under dark and red light (40 µmol m^-2^ s^-1^). *A. fabrum* suspensions were pipetted on root segment bundles and incubated for 2 days. The number of tumors was counted after 2 weeks. Mean of 3 experiments ± SE. (B) Results of *A. fabrum* root infection assay.

Stem infection assays were performed with *Nicotiana benthaminiana* plants that are larger and can be handled more easily than *Arabidopsis* plants (***Figure 6A***). *A. fabrum* cells were transferred to 2 injured sections on the same plant above each other. For a dark control, one section was covered with aluminum foil. Plants were kept for 1 d in red light, thereafter the aluminum foil was removed and bacterial cells were destroyed. After two weeks the formation of tumors could be observed in the dark controls of wild type or *agp1*^*-*^ infected stems, but not in the parts of the stems that were exposed to red light during infection. The results were reversed for the *agp2*^*-*^ mutant; in this case larger tumors were formed in the light and weak tumors in darkness. When the *agp1*^-^ / *agp2*^-^ double knockout was used for stem infection, there were no tumors or only small tumors formed both in light and darkness. After plant growth of 6 weeks, tumor sizes increased but the differences between the treatments remained (***Figure 6A***). These experiments were performed three times with qualitatively the same outcome.

**Figure 6.**
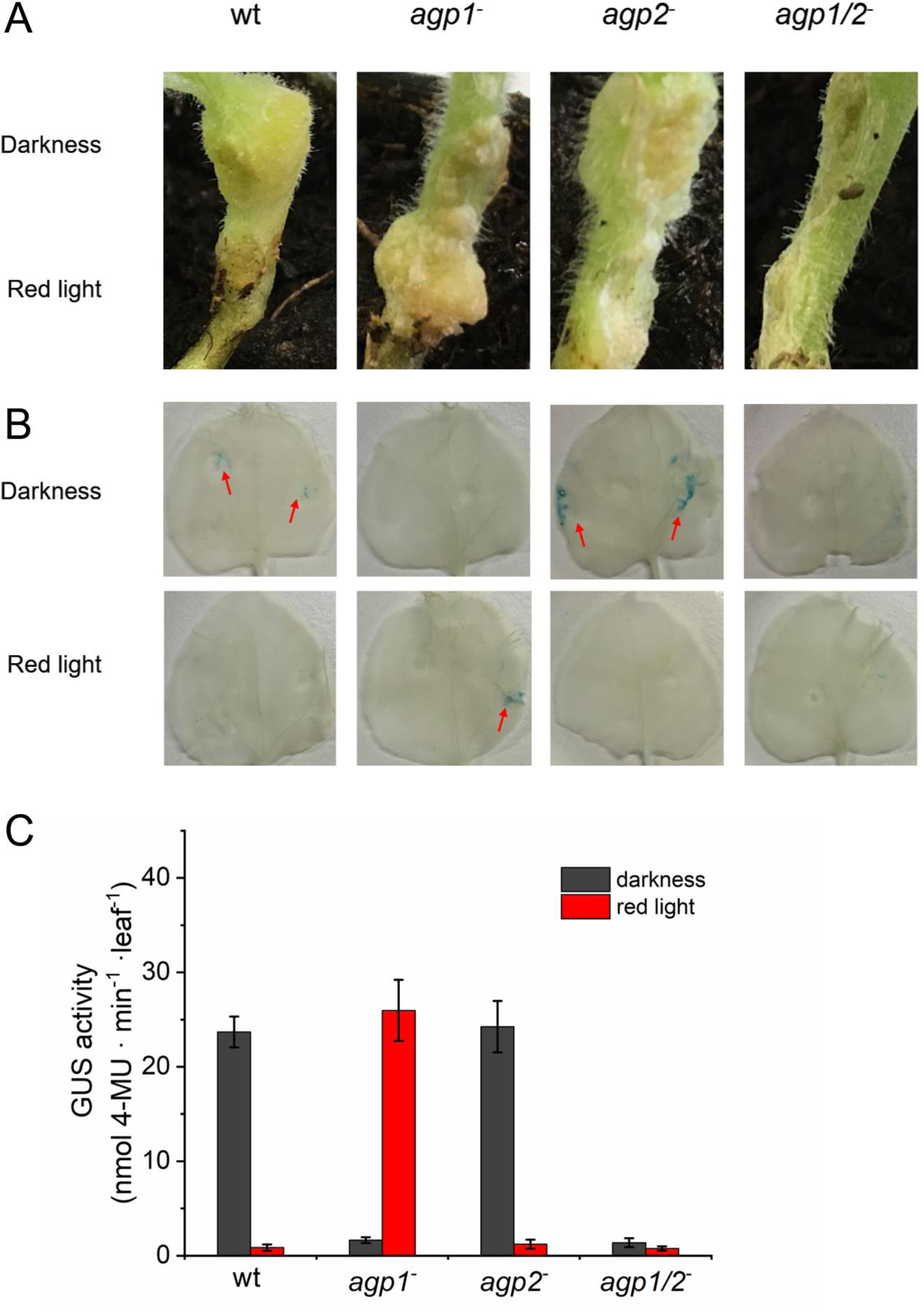
(A) Infection of *Nicotiana benthaminiana* stems by *A. fabrum* wild type (WT) and phytochrome mutants as indicated above the panels. During 1 d infection, the upper part of the stem was covered with aluminum and the entire plant placed in red light (1 µmol m^-2^ s^-1^). Stems with or without tumors were photographed after 6 weeks. The experiments were repeated 3 times with qualitatively identical outcome. (B) *Nicotiana benthaminiana* leaves were infected with *A. fabrum* WT and phytochrome mutants (*agp1*^*-*^, *agp2*^*-*^ and *agp1*^-^ / *agp2*^-^) and the GUS activity visualized as blue stains after treatment with X-Gluc. (C) Quantification of leaf infection assay by the MUG assay. Mean values of 3 independent infected leaves ± SE.

For quantification of infection we then performed a *Nicotiana benthaminiana* leaf infection assays using *A. fabrum* with the pBINGUS vector (15, 32). Following gene transfer from bacterium to plant cell, β-glucuronidase is expressed in the infected cells (***Figure 6C***). The expression of β-glucuronidase (GUS) can be visualized by enzymatic hydrolysis of X-Gluc which finally results in a blue staining of the infected cells. When infection was performed with type *A. fabrum* in the dark, blue colored spots appeared at several positions within the leaf. When the incubation was performed in red light, no blue spots appeared. When leaves were infiltrated with *agp1*^-^ / *agp2*^-^ double knockout mutant and kept in the dark or irradiated with red, staining was observed in neither case. Infiltration with *agp1*^-^ knockout mutant resulted in staining in the red irradiated leaf, but not in the dark control. In the *agp2*^-^ knockout mutant, the result was again reversed (or comparable to wild type): cells were stained in the dark sample, but not after the red light treatment. For quantification, the β-glucuronidase assay was then performed with 4-methylumbelliferyl-β-D-glucuronide (MUG) (33). After treating the leaves as above, leaf extracts were mixed with MUG and the fluorescence of the GUS product 4-methylumbelliferol (MU) was measured. For wild type *A. fabrum*, the MU signal was strong in the dark control, whereas red light treatment resulted in very low MU levels. With the *agp1*^-^ / *agp2*^*-*^ double knockout the fluorescence signal was always low. With the *agp1*^*-*^ knockout mutant, fluorescence was high upon red treatment but low in darkness. The light / dark pattern of the *agp2*^*-*^ knockout mutant was comparable to that of wild type *A. fabrum*.

All infection assays suggest that the gene transfer from *A. fabrum* to the plant is controlled by light and by both phytochromes. We would like to stress again that the loss of light regulation in the double knockout mutant implies that there is no impact of plant phytochrome in the light responses with e.g. wild type *A. fabrum*. In the *Nicotiana plumbaginifolia* assays (***Figure 6A-C***), the infection with *agp2*^*-*^ resulted in similar light / dark patterns as with wild type *A. fabrum*. This implies that Agp1 is alone responsible for the light control of infection. However, loss of Agp1 did not result in a reduction of the light response but in a reversion (stronger infection in red, low in dark). One could imagine that Agp1 reverts the light response of Agp2. Since there is an interaction between both proteins at lease in *vitro* (34), several molecular scenarios seem possible to explain such an impact of one phytochrome on the other.

### Microarray studies

Is there a role of Agp1 and Agp2 on gene expression? Quantification of mRNA of *Agrobacterium* wild type and the *agp1*^-^ / *agp2*^-^ double knockout mutant in darkness and light was performed by two independent approaches (data shown in ***Table S1***). Only 4 genes termed atu1049, atu3178, atu3179, atu3181 showed a more than 2-fold change between dark and light in both assays. For all other genes the difference was either smaller or obtained in one of the assays only and was thus regarded as insignificant. Atu1049 is annotated as a hypothetical protein and was found involved in GABA responses. atu3178, atu3179, atu3181 belong to an operon of a Zn^2+^ ABC transporter (35). The fact that three out of four genes of the same operon appear light regulated underpins the reliability of the assays. In the double knockout mutant, no gene appeared light regulated when the same analysis was performed, whereas the same genes (and more) were identified when dark grown *agp1*^*-*^ */ agp2*^*-*^ mutant and wild type were compared. All 23 genes coding for conjugal transfer proteins (tra proteins and mobC) had light / dark mean values in the range of 0.58 to 1.16. Although this could be indicative of a weak overall down regulation of *tra* gene expression, it is unlikely that the conjugal transfer is regulated by transcriptional activity. Moreover, no evidence for differences between wild type and double knockout mutant of *tra* transcripts was found. All 40 virulence (*vir*) genes had light / dark ratios (of single values) between 0.5 and 1.3. These genes encode proteins required for excision of T-DNA, transport and the type IV secretion system that are also important for conjugation. Again, the mRNA levels of the *agp1*^*-*^ / *agp2*^*-*^ double knockout mutant genes were in the same range as those of the wild type. All in all, on the transcriptional level there seems to be no significant phytochrome effect on and light regulation of genes that are relevant for DNA transfer to other cells or genes that could be responsible for growth or motility.

### Proteome studies

In order to find light versus dark or *agp1*^-^ / *agp2*^-^ double knockout mutant versus wild type differences on the level of proteins, we performed a comparative TMT proteome analysis (36). In this approach, proteins from cell extracts are digested by trypsin and covalently labeled with tags of slightly different molecular mass to mark each specific sample (***Table S2***). After labeling, samples are mixed and subjected to LC-MS/MS. The quantity of each peptide relative to the same peptide of another sample can be gained from comparison in the same run. The analysis was performed with 3 independent extracts of 4 different samples, wild type, double knockout both in dark and light. Out of 5400 *A. fabrum* proteins, 2814 were identified. We considered protein ratios of < 0.67 or > 1.5 with t-test probabilities < 0.05 as significant. An overview of the differences is given in the Venn diagram in ***Figure 7***. Of the 2814 detected proteins, 422 proteins were either light regulated or affected by the knockout mutation. In the wild type, 24 proteins appeared light regulated (***Table S3*** and ***Table S4***). Ten of the 24 proteins showed light / dark differences in the wild type only, but not in the double mutant (***Table S3***). Seven light regulated proteins have functions in energy metabolism, three are ribosomal proteins, two are signal transduction proteins, one is a diguanylate phosphodiesterase and one a Ras family protein. On this level, no overlap with the physiological functions discussed above is apparent. Quite interestingly, 24 proteins were light regulated in the double knockout mutant and not in the wild type. These effects could also be mediated by the photolyase PhrB, which as noted above was shown to control physiological responses in *A. fabrum*.

**Figure 7.**
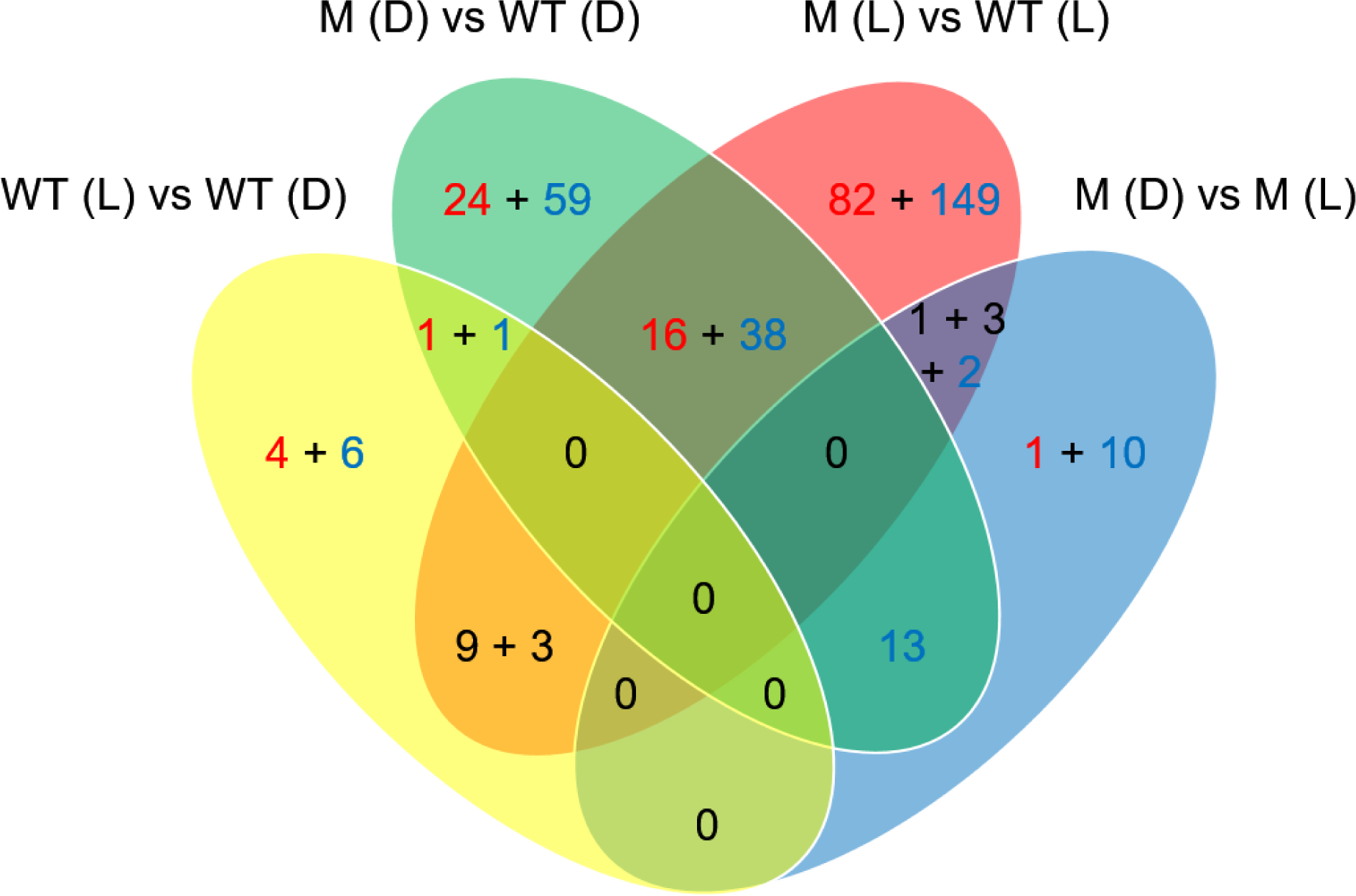
Differentially expressed proteins as identified by TMT analysis, Venn diagram. WT (D): wild type (darkness); WT (L): wild type (white light); M (D): mutant *agp1*^-^ / *agp2*^-^ (darkness); M (L): mutant *agp1*^-^ / *agp2*^-^ (white light). Ratio (experimental group vs control group) > 1.5 (*P* < 0.05) and < 0.67 (*P* < 0.05) were set as significantly up-regulated (red numbers) and down-regulated (blue numbers), respectively. The black numbers 9 + 3 indicate that 9 proteins were down-regulated in WT (L) vs. WT (D), and up-regulated in M (L) vs. WT (L) and 3 proteins were up-regulated in WT (L) vs. WT (D) and down-regulated in M (L) vs. WT (L). The black numbers 1 + 3 indicate that 1 protein was up-regulated in M (L) vs. WT (L) and down-regulated in M (D) vs. M (L) and 3 proteins were down-regulated in WT (L) vs. WT (L) and up-regulated in M (D) vs. M (L). The sum of differentially expressed proteins and identified protein was 422 and 2812, respectively. 134 and 353 proteins were only observed in both WT (D) and M (D) and both WT (L) and M (L), respectively.

More differences than between the light and dark grown cultures were found between the mutant and wild type strains. Altogether, 382 proteins had a different abundance in mutant and wild type (***Figure 7***). These results suggest that besides their roles as photoreceptors, phytochromes have also other functions both in darkness and in light, in line with the above phytochrome effects on motility and growth.

In the following, we focus on proteins related to motility, conjugation, plant infection and type IV secretion, for which phytochrome effects have been found.

#### Motility

Out of 8 detected chemotaxis proteins, one (McpC) showed a 0.4 fold ratio between mutant and wild type, whereas the others were not affected by light or mutant (***Table S5***). Of the 34 flagella proteins, 11 were detected in the assay (***Table S6***). For 8 of these, no significant differences between dark vs. light or mutant vs. wild type were detected. For FlaA and FlaB, the major constituents of the bacterial flagella, the protein levels in the mutant were ca. 1.7 fold higher as compared to the wild type.

The type IV pili might contribute to movement on agar surface. In the assay, 5 out of 10 type IV pili proteins were detected (***Table S7***). Two of these, CtpA and CtpE, revealed differences between mutant and wild type. CtpA of light samples was ca. 1.5 fold higher in the mutant, CtpE levels were 0.5 times lower in the mutant, both in light and dark samples.

#### Conjugation

Four of 22 proteins that are related to bacterial conjugation were detected in the assay (***Table S8***). All four proteins revealed differences between mutant and wild type. The central conjugation protein TraA cleaves the plasmid DNA, forms a covalent link with the DNA and unwinds the double stranded DNA. *A. fabrum* has three TraA proteins that are encoded on the circular chromosome, the AT-plasmid and the Ti-plasmid. Of these, only the AT-plasmid encoded TraA (Atu5111) was detected and the detection was only possible for dark samples. The levels of the mutant were about 3 times lower than of the wild type.

#### Virulence

Of the 26 proteins assigned to plant infection or virulence, only 2 were detected (***Table S9***), VirH1 and AcvB. Levels of both proteins were unchanged in mutant vs wild type or dark vs light.

#### Type IV secretion system

The type IV secretion system is important for both conjugation and plant infection. Eight of 26 proteins were detected (***Table S10***). The level of AvhB1 was different between wild type in the dark and wild type in the light, and for three others, AvhB4, AvhB9 and AvhB10, the levels were different between mutant and wild type.

#### Type VI secretion system

Through the type VI secretion system, toxin proteins are injected into competing bacterial cells (37). We found two proteins with wild type vs. mutant differences (***Table S11***) that belong to the type VI secretion system. The levels of the secretion protein Hcp were lower in mutant than in wild type cells (light and dark). One toxin protein, Atu4347, annotated as peptidoglycan amidase, had lower levels in the mutant (light) as compared to the wild type (light).

### Bacterial competition

These latter results prompted us to investigate the role of light and phytochromes in bacterial competition. In the first assay, we mixed *A. fabrum* with *E. coli* cells and measured the survival of *E.coli* by agar plate assays after appropriate dilution, similar as in (37). In this test we found no impact of *A. fabrum* wild type on the survival rate of *E. coli*, irrespective of whether the samples were kept in darkness, red light or far-red (***Figure 8A***). However, when the *agp1*^*-*^ knockout mutant was used for the competition assay, the survival of *E. coli* was drastically reduced. In the dark sample, only about 10% *E. coli* survived, and in red or far-red treated samples, survival was about one third of in the control. The *agp2*^*-*^ knockout mutant revealed wild type-like survival in dark and far-red treated samples, but reduced survival upon red light treatment. For the *agp1*^*-*^ */ agp2*^*-*^ double knockout mutant the values were similar to the wild type.

**Figure 8.**
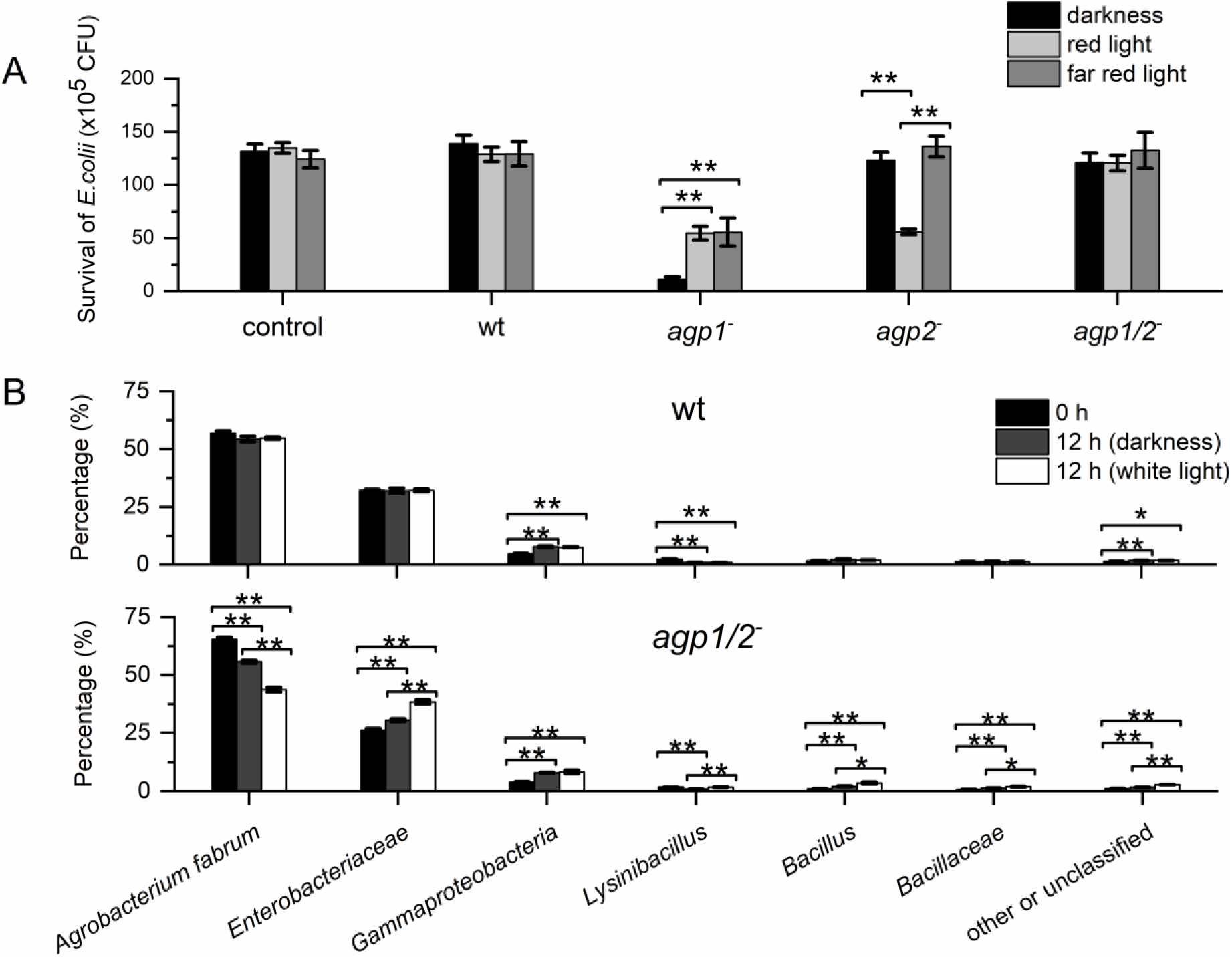
Competition assays of *A. fabrum* wild type and phytochrome mutant in darkness and light with *E. coli* (A) and with soil bacteria (B). In (A), *A. fabrum* was mixed with *E. coli* and the relative survival rate of *E. coli* determined after overnight culture. Mean values of 3 and 4 biological replicates ± SE. In (B), *A. fabrum* was mixed with soil bacteria and the relative content of each species determined by mass sequencing. Mean values of 4 biological replicates ± SE; ** = *P* <0.01; * = *P*<0.05.

We also performed a competition assay in which *A. fabrum* was mixed with bacteria from forest soil (***Figure 8B***). Here, quantification of the number of species was performed by next generation sequencing and the 16S rRNA sequence was used for species identification. When wild type *A. fabrum* was mixed with soil bacteria, the initial relative fraction of *A. fabrum* remained unchanged after 12 h co-incubation (***Figure 8B*** upper panel). When the *agp1*^-^ */ agp2*^-^ double mutant was used, the initial 62% fraction was reduced to 50% in darkness and even more reduced to 42% in white light (***Figure 8B*** lower panel). Enterobacteriaceae formed the largest group among the other bacteria, their fraction increased from 20% at the time of mixing to 30 or 35% upon 12 h co-cultivation in darkness or white light, respectively. We distinguished 5 other bacterial groups, and their light / dark patterns were similar to that of Enterobacteriaceae. The light effect observed in the experiment with the double knockout mutant could either be mediated through PhrB or could result from a light effect on the other soil bacteria. Because bacterial groups had higher levels after light treatment as compared to dark, the light effect on other bacteria seems unlikely. Although in the wild type, no light action is observed, the results of the double knockout mutant show that there must be a phytochrome mediated compensatory effect in the wild type.

## Discussion

We provide evidence for the involvement of bacterial phytochromes in regulation of growth, motility, plant infection, and interbacterial competition of *Agrobacterium fabrum* C58. The regulation of bacterial conjugation by phytochromes has been reported in an earlier study (15). Our proteome studies provide further information on phytochrome effects in *A. fabrum.* Since other *Agrobacterium* species have always Agp1 and/or Agp2 phytochrome homologs (***Figure 1***), we can assume that the modes of regulation are comparable in all *Agrobacterium* species.

The phytochrome-mediated regulation of plant infection can be regarded as the most important finding of the present work. The mechanism of DNA transfer has been studied intensely and is used by many botanical groups for plant transformation (38). All organs of plants contain plant phytochromes and other photoreceptors (39). We nevertheless assume that in our experiments the light effects on plant infection are mediated through the *A. fabrum* phytochromes, because the single knockouts always had a clear – although different – effect on the result and because with the *agp1*^-^ */ agp2*^-^ double knockout of *A. fabrum* there is no infection and no light response (***Figure 5*** and ***Figure 6***). A plant phytochrome effect would be more or less independent of the bacterial phytochrome situation. That all three effects – root, stem and leaf infection - have a similar pattern with respect to light and darkness and the response in wild type and mutants makes us confident that we observe true *A. fabrum* phytochrome effects. We assume that even in other work on plant transformation by *Agrobacterium*, where the impact of *Agrobacterium* on light regulation was not tested, the effect of light on transformation could have been mediated through *Agrobacterium* phytochromes, even though light sometimes had a positive (16) and sometimes a negative effect (17) on transformation. Engineering the phytochrome system of *A. fabrum* could lead to light independent transformation and increased transformation efficiencies.

What could be the evolutionary advantage of a phytochrome regulation for this unique gene transfer process? Light penetrates several millimeters into the soil, and far red light penetrates deeper than light of shorter wavelengths (40). Vessels of plant stems and roots guide sunlight deeper into the soil (41, 42). This light would inhibit plant transformation in the upper region of the soil during daytime. The reason for this inhibition could be that the plant releases enough exudates into the upper part of the soil during daytime so that *A. fabrum* would not benefit from tumor induction. Conjugation under natural condition results in a broader distribution of the Ti-plasmid (or AT plasmid) in the population of *A. fabrum* cells. Inhibition of conjugation leads thus to a lowering of plant transformation rates. Therefore, light and phytochrome effects on conjugation and transformation reinforce each other.

We found in *Nicotiana benthaminiana* transformation experiments that the *agp1*^*-*^ knockout mutant reverted the light / dark effect. With this mutant, plant transformation was high in the light and low in the dark (***Figure 6***). Both *A. fabrum* phytochromes interact in *vitro* (34), so Agp1 could act on Agp2 either by modulating other protein interactions or by covalent modifications. Phylogenetically, Agp1 and Agp2 are rather unrelated phytochromes. Agp2 has many homologs in related rhizobia. Among 22 *Agrobacterium* species, Agp2 homologs can be found in 21 species, whereas Agp1 homologs are present in only 4 of the species (***Figure 1***). In evolutionary terms, Agp2 is apparently present in *Agrobacterium* predecessors since a long time, whereas Agp1 was probably taken up later by horizontal gene transfer. In the time before the uptake of Agp1, light regulation of transformation was probably reverse, as may be tested by corresponding studies on *Agrobacterium* species that lack an Agp1 homolog.

The patterns of *A. fabrum* regulations with respect to light vs. dark and mutant effects differ largely. In principle, we can distinguish between “light-dependent” and “light-independent” phytochrome effects. (All effects are summarized in ***Table 1.***) Conjugation and plant infection are clearly light dependent. Both are connected with DNA transfer in which the type IV secretion system is involved. Both kinds of DNA transfer share further common features: plasmid DNA is nicked either by VirD2 or by TraA at a defined position, a single strand is covalently bound to the protein, DNA is unwound in a helicase reaction and the single stranded DNA-protein complex transported into the target cell. Quite interestingly, for each one of 4 the conjugation proteins that were detected in the TMT assay the levels were different between wild type and double knockout mutant. In case of the type IV secretion system, the same applies to 3 out of 8 proteins. Regulation of protein concentration by differential degradation could be one mechanism in the phytochrome regulation of DNA transfer. In mRNA microarrays there was no evidence for light or phytochrome regulation of transcription of any of the encoding genes. Since the differences in protein abundance between wild type and mutant are not large, and there was no clear light effect on protein levels, we consider, however, a different mechanism of signal transmission of light regulation. In conjugation and plant infection, dark action of phytochrome is reduced in the light. This pattern correlates with the histidine autokinase activity of Agp1 (3), but not with that of Agp2 (34). Phosphotransfer is thus a possible initial signal transmission mechanism, but cannot be the only one. Conjugation and plant infection could be mediated through an interaction of Agp1 and Agp2 with VirD2 and TraA, and could modulate the nuclease activities of the enzymes in a light dependent manner.

**Table 1.** Summary of phytochrome response in *A. fabrum*. The situation of wild type in darkness is regarded as normal. n.d. = not determined

|  | wild type |  | *agp1*<sup>-</sup> mutant |  | *agp2*<sup>-</sup> mutant |  | *agp1/2*<sup>-</sup> double knockout mutant |  |
| --- | --- | --- | --- | --- | --- | --- | --- | --- |
|  | dark | light | dark | light | dark | light | dark | light |
| **light regulated phytochrome effects** |  |  |  |  |  |  |  |  |
| conjugation | normal | low | low | very low | low | very low | zero | zero * |
| root infection | normal | low | n.d. | n.d. | n.d. | n.d. | low | very low |
| stem infection | normal | low | low | Normal | normal | low | very low | very low |
| leaf infection | normal | very low | very low | Normal | normal | very low | very low | very low |
| **not light regulated phytochrome effects** |  |  |  |  |  |  |  |  |
| motility at 26 °C | normal | normal | normal | lower | normal | lower | normal | normal ** |
| motility at 37 °C | normal | normal | normal | dead at extreme pH | normal | dead at extreme pH | normal | normal ** |
| growth at 28 °C | normal | normal | high | high | high | high | high | high |
| growth at 37 °C | normal | normal | normal | normal | low | normal | low | low |
| interbacterial competition | normal | normal | strong | intermediate | normal | intermediate | normal | normal |

\* (15)
\*\* simplified from results at different pH

As “light independent” effects we consider effects where no light-dark differences were found in the wild type under standard conditions, but where differences between mutant and wild type were found. Effects on growth at ambient temperature and effects on interbacterial competition belong to this category (***Table 1***). We would also assign effects on motility to this category, because in the wild type there is no light regulation; this is only uncovered in single knockout mutants. In addition to mutant vs. wild type differences, light effects were observed under special conditions in these otherwise light independent phytochrome responses. In motility assays, an inhibitory red light effect was found in single knockout mutants at pH 5 (***Figure 4H***). In cell growth, light effects were not observed at 28 °C but at 37 °C (***Figure 2***). In the proteome studies (***Figure 7***), the levels of more proteins differed between double knockout mutant than there were differences and wild type imply that there are more effects in *A. fabrum* that belong to this category of light-independent effects.

It is possible that light regulation provides evolutionary advantages only under some conditions and not under others. In such a case, an effect could appear unregulated by light under standard conditions but still be influenced by phytochrome, as we observed for some light-independent effects. Alternatively, an effect could have been under light control in an evolutionary progenitor of *A. fabrum*. With the adaptation to the soil environment, light regulation of this effect could have been lost. Despite their reduced roles in the soil, phytochromes “survived” because they were still required for the light dependent phytochrome effects. Clearly, the loss of a phytochrome (in a mutant) would result in a loss of light regulation, but it could also result in a distortion of other effects that were only previously under light control, the light-independent effects.

The present study, together with earlier work, shows that both DNA transfer processes of *A. fabrum*, conjugation and plant infection, are controlled by light and by phytochromes. Based on microarray and proteome studies, we exclude differential gene activation as a first step of phytochrome regulation and propose a direct modulation of TraA or VirD2, proteins that catalyze the first steps in DNA transfer processes. Effects on growth, interbacterial competition and motility are described as light independent in the wild type. An impact of phytochrome and light is uncovered by knockout mutants and / or elevated temperature.

## Materials and methods

See supplemental material

## Acknowledgments

The project was supported by DFG grants LA 799/7-3 and KR 2034/1 to Tilman Lamparter and Norbert Krauß. Patrick Scheerer was supported by the DFG: SFB1078-B06, SFB1365-A03 and DFG under Germany’s Excellence Strategy—EXC 311 2008/1 (UniSysCat)—390540038 (Research Unit E). Peng Xue, Yingnan Bai and Yuanyuan Ma were supported by the China Scholarship Council.

## Additional information

Funding

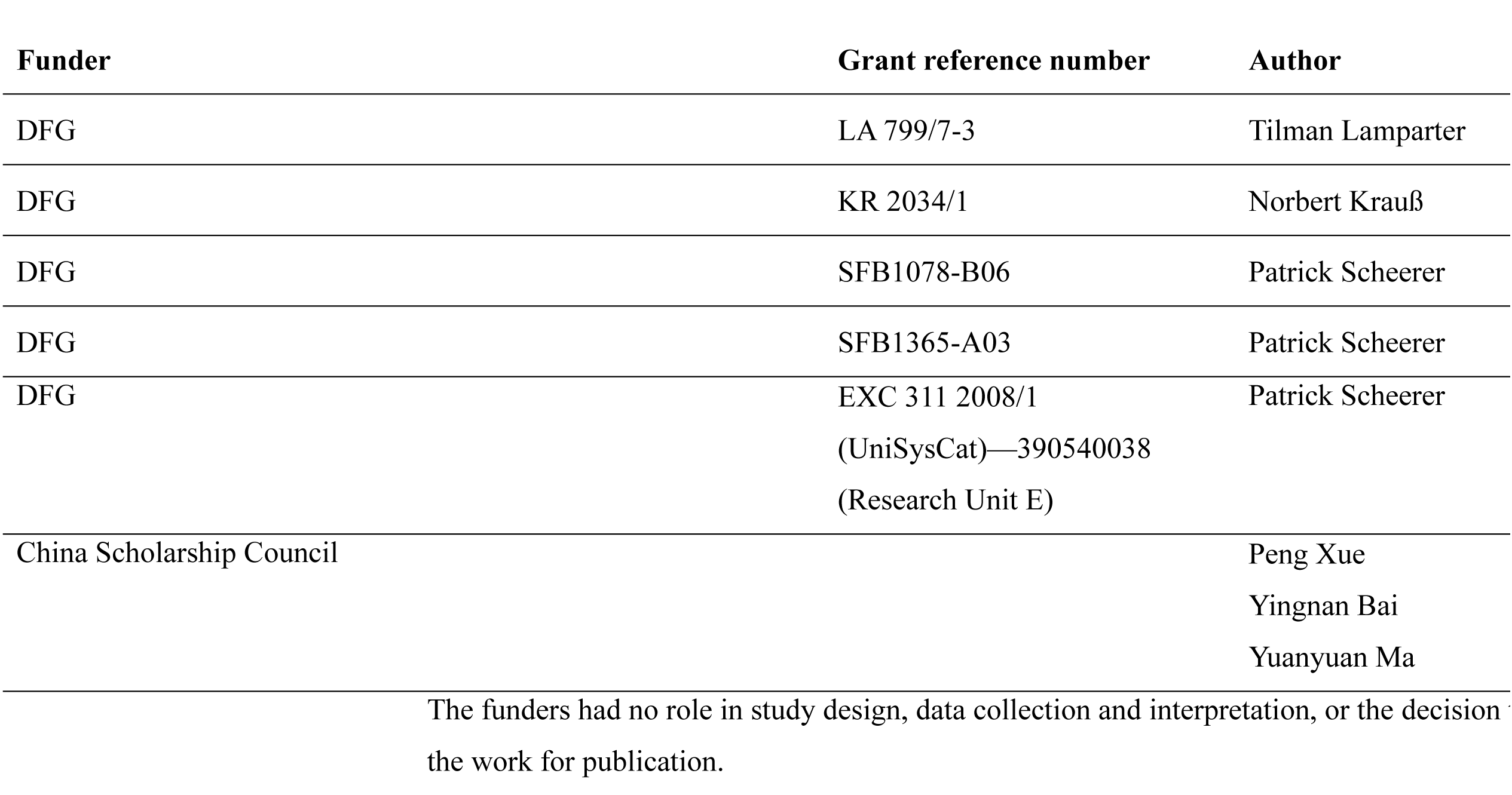

### Author contributions

Tilman Lamparter and Peng Xue designed the experiments, analyzed the data and wrote the manuscript with input from Norbert Krauß and Patrick Scheerer;. Yingnan Bai performed cell motility assays; Gregor Rottwinkel performed root infection assays and contributed to microarray assays; Elizaveta Averbukh contributed to *Nicotiana benthamiana* infection; Yuanyuan Ma and Thomas Roeder contributed to bacterial competition experiments; Peng Xue performed the other experiments. All authors reviewed the manuscript.

## Additional files

### Supplementary Tables

**Supplementary Table S1 is separate**

**Supplementary Table S2.** Sample labeling information. D: darkness; L: white light; WT_D: wild type_darkness; WT_L: wild type_white light; M_D: mutant_darkness; M_L: mutant_white light

| Group ID | Sample Name | Label Reagent |
| --- | --- | --- |
| D | M_D1 | TMT10-126 |
| D | M_D2 | TMT10-127N |
| D | M_D3 | TMT10-127C |
| D | WT_D1 | TMT10-128N |
| D | WT_D2 | TMT10-128C |
| D | WT_D3 | TMT10-130N |
| L | M_L1 | TMT10-126 |
| L | M_L2 | TMT10-129N |
| L | M_L3 | TMT10-129C |
| L | WT_L1 | TMT10-130N |
| L | WT_L2 | TMT10-130C |
| L | WT_L3 | TMT10-131 |

**Supplementary Table S3.** iTRAQ/TMT assay, ratios between dark and light for 10 proteins which are significantly higher or lower in dark vs. light of wild type, no significant L/D difference in double knockout (see yellow field in Venn diagram ***Figure 7***)

| Protein | Function | Fold change >1.5 or < 0.67; *P* < 0.05 |
| --- | --- | --- |
| 6 proteins |  |  |
| Rho (Atu2833) | Transcription termination factor | WT(D) / WT(L)=1.56 |
| ClpP (Atu1258) | ATP-dependent Clp protease proteolytic subunit 2 | WT(D) / WT(L)=1.67 |
| NuoI (Atu1278) | NADH-quinone oxidoreductase subunit I | WT(D) / WT(L)=1.59 |
| RpsL (Atu1951) | 30S ribosomal protein S12 | WT(D) / WT(L)=1.82 |
| AtpI (Atu0713) | ATP synthase protein I | WT(D) / WT(L)=1.85 |
| Atu6048 | putative DEAD/DEAH box helicase | WT(D) / WT(L)=1.56 |
| 4 proteins |  |  |
| CobL (Atu2798) | Precorrin-6y methyltransferase | WT(D) / WT(L)=0.67 |
| Atu2197 | putative diguanylate phosphodiesterase | WT(D) / WT(L)=0.51 |
| Atu4876 | Putative oxidoreductase | WT(D) / WT(L)=0.57 |
| Atu6114 | putative Ras family protein | WT(D) / WT(L)=0.63 |

**Supplementary Table S4.**
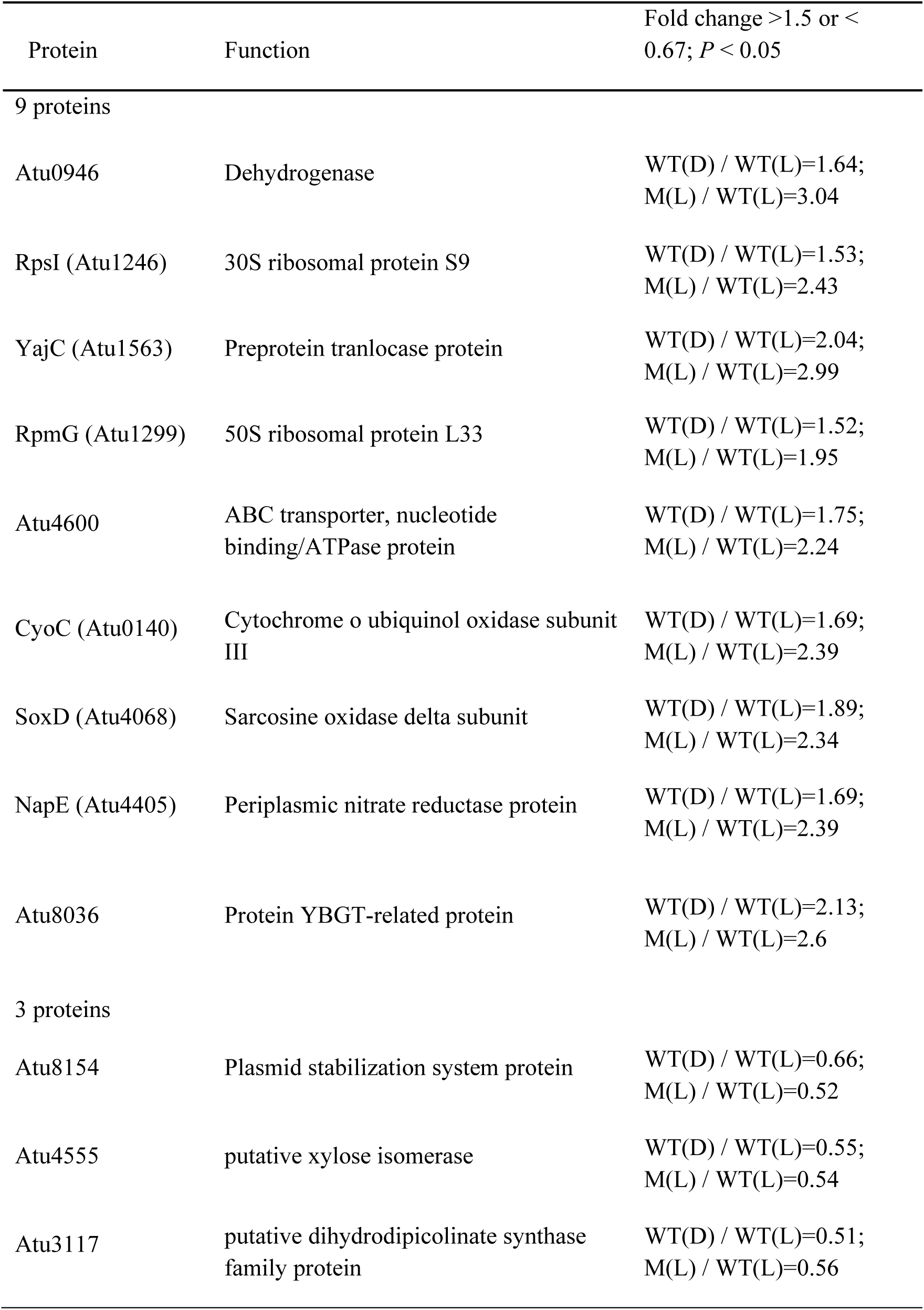
iTRAQ/TMT assay, ratios between dark and light for 13 proteins which are significantly higher or lower in dark vs. light of wild type and the *agp1*^-^ / *agp2*^-^ double knockout mutant. See yellow and red overlapping field in Venn diagram ***Figure 7***.

| Protein | Function | Fold change >1.5 or <0.67; *P* < 0.05 |
| --- | --- | --- |
| 9 proteins |  |  |
| Atu0946 | Dehydrogenase | WT(D) / WT(L)=1.64;<br>M(L) / WT(L)=3.04 |
| RpsI (Atu1246) | 30S ribosomal protein S9 | WT(D) / WT(L)=1.53;<br>M(L) / WT(L)=2.43 |
| YajC (Atu1563) | Preprotein translocase protein | WT(D) / WT(L)=2.04;<br>M(L) / WT(L)=2.99 |
| RpmG (Atu1299) | 50S ribosomal protein L33 | WT(D) / WT(L)=1.52;<br>M(L) / WT(L)=1.95 |
| Atu4600 | ABC transporter, nucleotide binding/ATPase protein | WT(D) / WT(L)=1.75;<br>M(L) / WT(L)=2.24 |
| CyoC (Atu0140) | Cytochrome o ubiquinol oxidase subunit III | WT(D) / WT(L)=1.69;<br>M(L) / WT(L)=2.39 |
| SoxD (Atu4068) | Sarcosine oxidase delta subunit | WT(D) / WT(L)=1.89;<br>M(L) / WT(L)=2.34 |
| NapE (Atu4405) | Periplasmic nitrate reductase protein | WT(D) / WT(L)=1.69;<br>M(L) / WT(L)=2.39 |
| Atu8036 | Protein YBGT-related protein | WT(D) / WT(L)=2.13;<br>M(L) / WT(L)=2.6 |
| 3 proteins |  |  |
| Atu8154 | Plasmid stabilization system protein | WT(D) / WT(L)=0.66;<br>M(L) / WT(L)=0.52 |
| Atu4555 | putative xylose isomerase | WT(D) / WT(L)=0.55;<br>M(L) / WT(L)=0.54 |
| Atu3117 | putative dihydrodipicolinate synthase family protein | WT(D) / WT(L)=0.51;<br>M(L) / WT(L)=0.56 |

**Supplementary Table S5.** The list of detected proteins of chemotaxis proteins of flagellar system of bacterial motility proteins of *A. fabrum*. CheA (Atu0517), Atu0515, CheD (Atu0521), CheR (Atu0518), Atu4805, CheD (Atu2618), McpA (Atu3094), McpA (Atu6132), McpG (Atu0738), McpA (Atu2360), Atu0514, McpA (Atu2173), Atu0373, MclA (Atu1912), MclA (Atu0526), McpA (Atu0387), McpA (Atu2223), Atu5442, Atu4736 and Atu3725 were not detected. WT (D): wild type (darkness); WT (L): wild type (white light); M (D): mutant (darkness); M (L): mutant (white light); Mean values of 3 biological replicates ± SE.

| Protein | Abundances | | | | Fold change >1.5 or < 0.67;<br>$P < 0.1$ (*) or $P < 0.05$ (**) |
| --- | --- | --- | --- | --- | --- |
|  | WT (D) | WT (L) | M (D) | M (L) |  |
| CheB (Atu0519) | 117 ± 40 | 115 ± 15 | 106 ± 28 | 85 ± 8 | |
| CheW (Atu2075) | 114 ± 5 | | 100 ± 25 | | |
| CheW (Atu2617) | 99 ± 10 | 118 ± 8 | 84 ± 2 | 82 ± 10 | |
| CheY (Atu0516) | 95 ± 8 | 102 ± 6 | 112 ± 25 | 98 ± 10 | |
| CheY (Atu0520) | 114 ± 2 | 113 ± 11 | 80 ± 3 | 87 ± 8 | |
| McpA (Atu0646) | | 133 ± 36 | | 67 ± 10 | |
| McpC (Atu0872)<br>: methyl-<br>accepting<br>chemotaxis<br>protein | 118 ± 24 | | 47 ± 10 | | M(D) / WT(D)=0.40 * |
| McpV (Atu1027) | 83 ± 9 | 65 ± 6 | 112 ± 22 | 135 ± 57 | |

**Supplementary Table S6.**
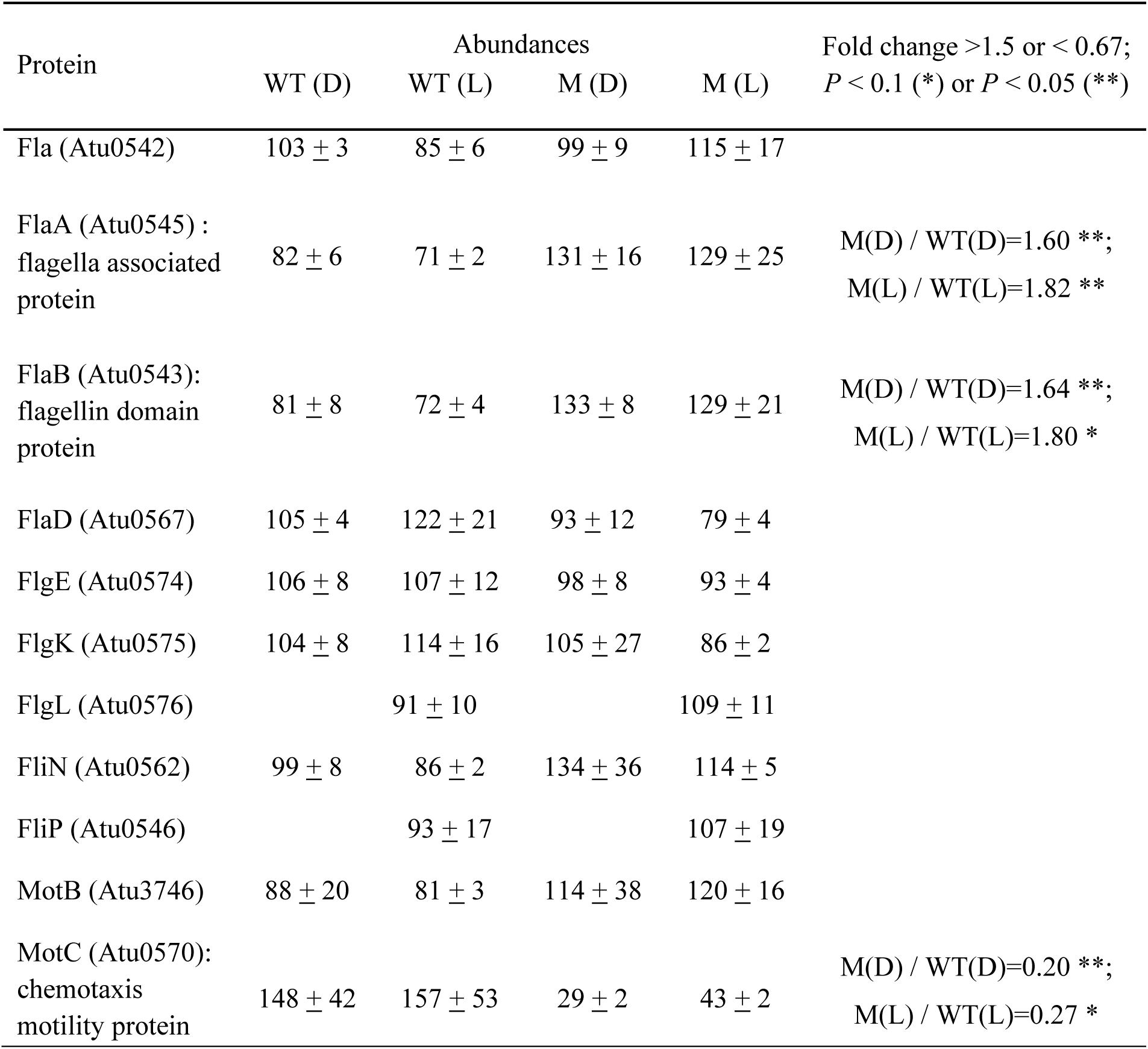
The list of detected proteins of flagellar assembly proteins of flagellar system of bacterial motility proteins of *A. fabrum*. FlaF (Atu0577), FlbT (Atu0578), FlgA (Atu0551), FlgB (Atu0555), FlgC (Atu0554), FlgD (Atu0579), FlgF (Atu0558), FlgG (Atu0552), FlgH (Atu0548), FlgI (Atu0550), FlhA (Atu0581), FlhB (Atu0564), FliE (Atu0553), FliF (Atu0523), FliG (Atu0563), FliI (Atu0557), FliL (Atu0547), FliM (Atu0561), FliQ (Atu0580), FliR (Atu0582), MotA (Atu0560), MotB (Atu0569) and MotD (Atu0571) were not detected. WT (D): wild type (darkness); WT (L): wild type (white light); M (D): mutant (darkness); M (L): mutant (white light); Mean values of 3 biological replicates ± SE.

| Protein | Abundances |  |  |  | Fold change >1.5 or < 0.67;<br>*P* < 0.1 (*) or *P* < 0.05 (**) |
| --- | --- | --- | --- | --- | --- |
|  | WT (D) | WT (L) | M (D) | M (L) |  |
| Fla (Atu0542) | 103 ± 3 | 85 ± 6 | 99 ± 9 | 115 ± 17 | |
| FlaA (Atu0545) :<br>flagella associated<br>protein | 82 ± 6 | 71 ± 2 | 131 ± 16 | 129 ± 25 | M(D) / WT(D)=1.60 **;<br>M(L) / WT(L)=1.82 ** |
| FlaB (Atu0543):<br>flagellin domain<br>protein | 81 ± 8 | 72 ± 4 | 133 ± 8 | 129 ± 21 | M(D) / WT(D)=1.64 **;<br>M(L) / WT(L)=1.80 * |
| FlaD (Atu0567) | 105 ± 4 | 122 ± 21 | 93 ± 12 | 79 ± 4 | |
| FlgE (Atu0574) | 106 ± 8 | 107 ± 12 | 98 ± 8 | 93 ± 4 | |
| FlgK (Atu0575) | 104 ± 8 | 114 ± 16 | 105 ± 27 | 86 ± 2 | |
| FlgL (Atu0576) | | 91 ± 10 | | 109 ± 11 | |
| FliN (Atu0562) | 99 ± 8 | 86 ± 2 | 134 ± 36 | 114 ± 5 | |
| FliP (Atu0546) | | 93 ± 17 | | 107 ± 19 | |
| MotB (Atu3746) | 88 ± 20 | 81 ± 3 | 114 ± 38 | 120 ± 16 | |
| MotC (Atu0570):<br>chemotaxis<br>motility protein | 148 ± 42 | 157 ± 53 | 29 ± 2 | 43 ± 2 | M(D) / WT(D)=0.20 **;<br>M(L) / WT(L)=0.27 * |

**Supplementary Table S7.** The list of detected proteins of pilus system of bacterial motility proteins of *A. fabrum*. PilA (Atu3514), CtpB (Atu0223), CtpF (Atu0219), CtpG (Atu0218) and Atu4731 were not detected. WT (D): wild type (darkness); WT (L): wild type (white light); M (D): mutant (darkness); M (L): mutant (white light); Mean values of 3 biological replicates ± SE.

| Protein | Abundances | | | | Fold change $>1.5$ or $<0.67$;<br>$P < 0.1$ (*) or $P < 0.05$ (**) |
| --- | --- | --- | --- | --- | --- |
|  | WT (D) | WT (L) | M (D) | M (L) |  |
| CtpA (Atu0224):<br>components of<br>type IV pilus,<br>pilin subunit | $94 \pm 5$ | $79 \pm 7$ | $111 \pm 33$ | $121 \pm 9$ | M(L) / WT(L)=1.53 ** |
| CtpC (Atu0222) | $82 \pm 20$ | $118 \pm 13$ | $78 \pm 6$ | $82 \pm 5$ | |
| CtpD (Atu0221) | $99 \pm 6$ | $97 \pm 6$ | $135 \pm 24$ | $103 \pm 10$ | |
| CtpE (Atu0220):<br>components of<br>type IV pilus | $175 \pm 23$ | $130 \pm 13$ | $58 \pm 15$ | $70 \pm 3$ | M (D) / WT (D)=0.33 **;<br>M (L) / WT (L)=0.54 ** |
| Atu4732 | $93 \pm 12$ | $99 \pm 4$ | $101 \pm 3$ | $101 \pm 14$ | |

**Supplementary Table S8.** The list of detected proteins of conjugation of *A. fabrum*. TraA (Atu4855), TraA (Atu6127), TraB (Atu6129), TraC (Atu5110), TraF (Atu6128), TraG (Atu5108), TraG (Atu6124), TraH (Atu6130), TrbB (Atu6041), TrbC (Atu6040), TrbD (Atu6039), TrbE (Atu6038), TrbF (Atu6034), TrbG (Atu6033), TrbH (Atu6032), TrbI (Atu6031), TrbJ (Atu6037), TrbL (Atu6035) were not detected. WT (D): wild type (darkness); WT (L): wild type (white light); M (D): mutant (darkness); M (L): mutant (white light); Mean values of 3 biological replicates ± SE.

| Protein | Abundances |  |  |  | Fold change >1.5 or < 0.67;<br>P < 0.1 (*) or P < 0.05 (**) |
| --- | --- | --- | --- | --- | --- |
|  | WT (D) | WT (L) | M (D) | M (L) |  |
| MobC (Atu4857):<br>mobilization<br>protein C | 87 ± 9 | 96 ± 6 | 138 ± 15 | 104 ± 11 | M (D) / WT (D)=1.59 ** |
| TraA (Atu5111):<br>conjugal transfer<br>relaxase | 147 ± 22 | | 49 ± 15 | | M (D) / WT (D)=0.33 ** |
| TraC (Atu6126):<br>conjugal transfer<br>protein | 69 ± 6 | 67 ± 6 | 161 ± 13 | 133 ± 13 | M (D) / WT (D)=2.34 **;<br>M (L) / WT (L)=2 ** |
| TraD (Atu5109):<br>conjugal transfer<br>protein | 85 ± 13 | 87 ± 1 | 133 ± 14 | 113 ± 5 | M (D) / WT (D)=1.56 * |

**Supplementary Table S9.**
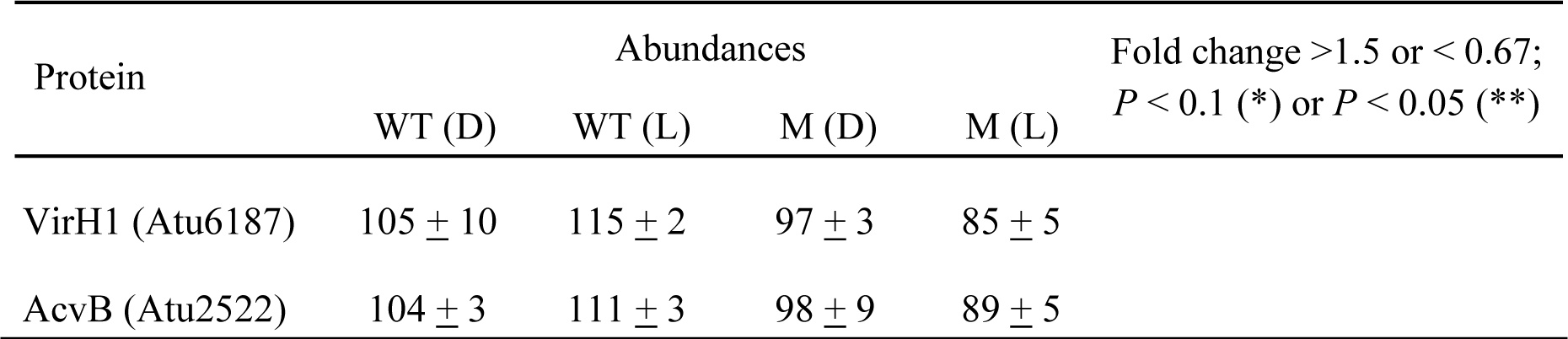
The list of detected proteins of virulence proteins without a protein of type IV secretion system of *A. fabrum*. VirA (Atu6166), VirC1 (Atu6180), VirC2 (Atu6179), VirD1 (Atu6181), VirD2 (Atu6182), VirD3 (Atu6183), AvhD4 (Atu4858), VirD4 (Atu6184), VirD5 (Atu6185), VirE0 (Atu6188), VirE1 (Atu6189), VirE2 (Atu6190), VirE3 (Atu6191), VirE3 (Atu6186), VirF (Atu6154), VirG (Atu6178), VirK (Atu6156), MviN (Atu0347), TrlR (Atu6192), Atu6193, Atu6194, Atu6195, Atu6196 and Atu6197 were not detected. WT (D): wild type (darkness); WT (L): wild type (white light); M (D): mutant (darkness); M (L): mutant (white light); Mean values of 3 biological replicates ± SE.

| Protein | Abundances |  |  |  | Fold change >1.5 or < 0.67;<br>P < 0.1 (*) or P < 0.05 (**) |
| --- | --- | --- | --- | --- | --- |
|  | WT (D) | WT (L) | M (D) | M (L) |  |

|  |  |  |  |  |
| --- | --- | --- | --- | --- |
| VirH1 (Atu6187) | 105 ± 10 | 115 ± 2 | 97 ± 3 | 85 ± 5 |
| AcvB (Atu2522) | 104 ± 3 | 111 ± 3 | 98 ± 9 | 89 ± 5 |

**Supplementary Table S10.** The list of detected proteins of type IV secretion system of *A. fabrum*. VirB1 (Atu6167), AvhB2 (Atu5163), VirB2 (Atu6168), AvhB3 (Atu5164), VirB3 (Atu6169), VirB4 (Atu6170), VirB5 (Atu6171), AvhB6 (Atu5167), VirB6 (Atu6172), VirB8 (Atu6174), VirB9 (Atu6175), VirB10 (Atu6176), AvhB11 (Atu5172), VirB11 (Atu6177), Atu4858 (traG), Atu5108 (traG), Atu6124 (traG) and virD4 (Atu6184) of type IV secretion system were not detected. WT (D): wild type (darkness); WT (L): wild type (white light); M (D): mutant (darkness); M (L): mutant (white light); Mean values of 3 biological replicates ± SE.

| Protein | Abundances |  |  |  | Fold change >1.5 or < 0.67;<br>P < 0.1 (*) or P < 0.05 (**) |
| --- | --- | --- | --- | --- | --- |
|  | WT (D) | WT (L) | M (D) | M (L) |  |
| AvhB1 (Atu5162):<br>type IV secretion<br>protein | 59 ± 4 | 88 ± 9 | 145 ± 19 | 112 ± 3 | WT (L) / WT (D)=1.50 **;<br>M (D) / WT (D)=2.46 ** |
| AvhB4 (Atu5165):<br>type IV secretion<br>protein | 78 ± 3 | 69 ± 5 | 133 ± 29 | 131 ± 23 | M (L) / WT (L)=1.89 * |
| AvhB5 (Atu5166) | 69 ± 1 | 100 ± 2 | 132 ± 57 | 100 ± 5 |  |
| AvhB7 (Atu5168):<br>type IV secretion<br>protein | 82 ± 11 | 79 ± 11 | 121 ± 23 | 121 ± 4 | M (L) / WT (L)=1.52 ** |
| VirB7 (Atu6173) | 100 ± 5 |  | 133 ± 21 |  |  |
| AvhB8 (Atu5169) | 89 ± 19 | 96 ± 13 | 100 ± 20 | 104 ± 9 |  |
| AvhB9 (Atu5170):<br>type IV secretion<br>protein | 74 ± 4 | 81 ± 9 | 137 ± 2 | 119 ± 7 | M (D) / WT (D)=1.85 ** |
| AvhB10<br>(Atu5171): type IV<br>secretion protein | 74 ± 5 | 66 ± 5 | 136 ± 1 | 134 ± 4 | M (D) / WT (D)=1.84 **;<br>M (L) / WT (L)=2.01 ** |

**Supplementary Table S11.**
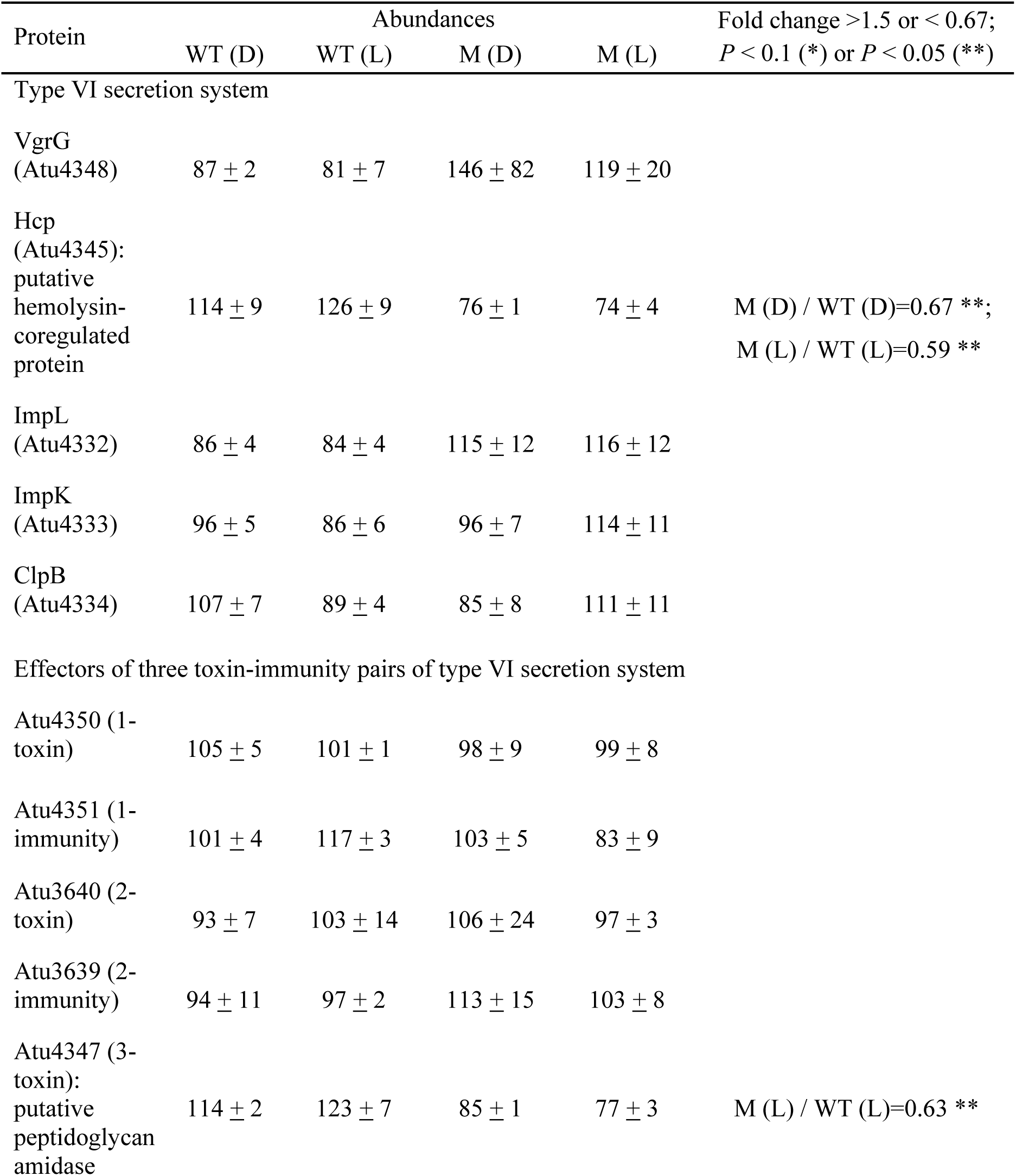
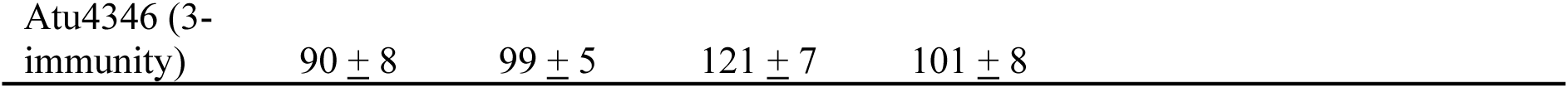
The list of detected proteins of Type VI secretion system and three toxin-immunity pairs – *A. fabrum*. WT (D): wild type (darkness); WT (L): wild type (white light); M (D): mutant (darkness); M (L): mutant (white light); Mean values of 3 biological replicates ± SE.

